# The proteomic landscape of proteotoxic stress in a fibrogenic liver disease

**DOI:** 10.1101/2024.11.01.621457

**Authors:** Florian A. Rosenberger, Sophia C. Mädler, Katrine Holtz Thorhauge, Sophia Steigerwald, Malin Fromme, Mikhail Lebedev, Caroline A. M. Weiss, Marc Oeller, Maria Wahle, Maximilian Zwiebel, Niklas A. Schmacke, Sönke Detlefsen, Peter Boor, Joseph Kaserman, Andrew Wilson, Ondřej Fabián, Soňa Fraňková, Aleksander A. Krag, Pavel Strnad, Matthias Mann

## Abstract

Protein misfolding diseases, including alpha-1 antitrypsin deficiency (AATD), pose significant health challenges, with their cellular progression still poorly understood^1–3^. We utilize spatial proteomics by mass spectrometry and machine learning to map AATD in human liver tissue. Combining Deep Visual Proteomics (DVP) with single-cell analysis^4,5^, we probe intact patient biopsies to resolve molecular events during hepatocyte stress in pseudo-time across fibrosis stages. We achieve unprecedented proteome depth of up to 3,800 proteins from a third of a single cell in formalin-fixed, paraffin-embedded (FFPE) tissue. This dataset revealed a potentially clinically actionable peroxisomal upregulation that precedes the canonical unfolded protein response. Our single-cell proteomics data show alpha-1 antitrypsin accumulation is largely cell-intrinsic, with minimal stress propagation between hepatocytes. We integrated proteomic data with AI-guided image-based phenotyping across multiple disease stages, revealing a terminal hepatocyte state characterized by globular protein aggregates and distinct proteomic signatures, notably including elevated TNFSF10/TRAIL expression. This phenotype may represent a critical disease progression stage. Our study offers novel insights into AATD pathogenesis and introduces a powerful methodology for high-resolution, in situ proteomic analysis of complex tissues. This approach holds potential to unravel molecular mechanisms in various protein misfolding disorders, setting a new standard for understanding disease progression at the single-cell level in human tissue.

## Introduction

Spatial omics technologies are revolutionizing our ability to deconvolute molecular events at single-cell resolution within a tissue context. While much focus has been placed on spatial genomics and transcriptomics, recent advances in multiplexed imaging and proteomics are beginning to shed light on the functional proteomic layer. Mass spectrometry-based proteomics has made significant strides towards biologically informative single-cell analysis, now enabling quantification of up to 5,000 proteins in cultured cells^6,7^. In the tissue context, we have recently introduced Deep Visual Proteomics (DVP), which integrates staining, AI-guided cell segmentation and classification, laser microdissection of single-cell shapes, and high-sensitivity mass spectrometry^4,5^. DVP excels in digital pathology applications with pronounced spatial and visual components, providing simultaneous and deep proteomic characterization at the level of thousands of proteins.

We reasoned that these emerging technologies would be ideally suited to elucidate molecular events during the progressive worsening of proteotoxicity as it unfolds in patients. Proteotoxicity, characterized by the accumulation of misfolded and aggregated proteins leading to cell damage, is a hallmark of many diseases, including neurodegenerative pathologies such as Alzheimer’s and Parkinson’s disease^8–10^. The underlying cause of proteotoxicity is a disruption in protein homeostasis, resulting in an imbalance between protein synthesis, folding, and clearance mechanisms^3^.

To investigate proteotoxicity in a clinically relevant context, we focused on a disorder with unmet clinical need that exemplifies the challenges of protein misfolding and aggregation in a vital organ. The fibrogenic liver disease alpha-1 antitrypsin deficiency (AATD), is a genetic disorder caused by autosomal, co-dominant mutations in the *SERPINA1* gene resulting in misfolding and accumulation of alpha-1 antitrypsin (AAT) in hepatocytes. Most severe AATD cases are caused by a homozygous Z-variant (Pi*ZZ genotype) with a peak incidence of 1:2,000 in individuals of European descent^1,2,11,12^. Current hypotheses suggest that the severity of liver damage correlates with the amount of accumulated AAT^13–18^. However, the mechanisms driving fibrogenesis or hepatocyte survival versus death remain unclear, leaving potentially druggable targets unexplored.

## Results

To address this challenge, we curated a cohort of formalin-fixed paraffin-embedded (FFPE) biopsies and liver explants from patients homozygous for the pathogenic Z-variant (Pi*ZZ), encompassing all fibrosis stages (n = 35, Extended Data Fig. 1a). Despite the same underlying disease-causing mutation at a similar median age (57.3 ± SD 9.9 years) and BMI (25.4 ± SD 4.0), the fibrosis stages varied drastically, indicating unexplored molecular resilience or risk profiles.

### Proteomic mapping of hepatocyte responses to proteotoxic stress

To elucidate the molecular basis of the observed clinical heterogeneity in AATD patients, we implemented a comprehensive proteomic mapping approach to characterize hepatocyte responses to proteotoxic stress. We first laser microdissected 3 µm thick FFPE sections from patient biopsies and analyzed them with mass spectrometry following our DVP workflow. After staining for cell outlines and AAT, we segmented and stratified cells into low, moderate, and high aggregate load groups based on their microscopy images (Fig. 1a and 1b). The proteome of 100 shapes, equivalent to the volume of 10–15 complete hepatocytes, was then acquired on the recently introduced Orbitrap Astral mass spectrometer, yielding a high-quality dataset with a mean proteomic depth exceeding 5,000 proteins per sample (Extended Data Fig. 1b and 1c). We observed a striking 32-fold difference in AAT levels between low and high-load cells. The AAT load was captured on the second principal component, preceded only by the fibrosis stage on the first component (Extended Data Fig. 1d to 1f). Given the sparsity of AAT+ cells in biopsy material, this validated our laser microdissection approach as it allowed the biological phenotype to emerge more clearly. Biopsies with a low fibrosis stage exhibited lower AAT baseline loading compared to high fibrosis stages on both proteomics and imaging data, while the maximum load remained fairly equal across all stages (Extended Data Fig. 1g). The proteomes of the three load classes differed markedly (16.2% significant hits at < 5% FDR, paired two-sided t-test; Fig. 1c). Alongside AAT, several known markers of AATD liver pathology were highly enriched in aggregate-positive cells, such as a 1.8-fold increased ER chaperone HSPA5 and a 2.8-fold increased ER-Golgi cargo receptor LMAN1 (Fig. 1d)^19–21^.

**Figure 1:**
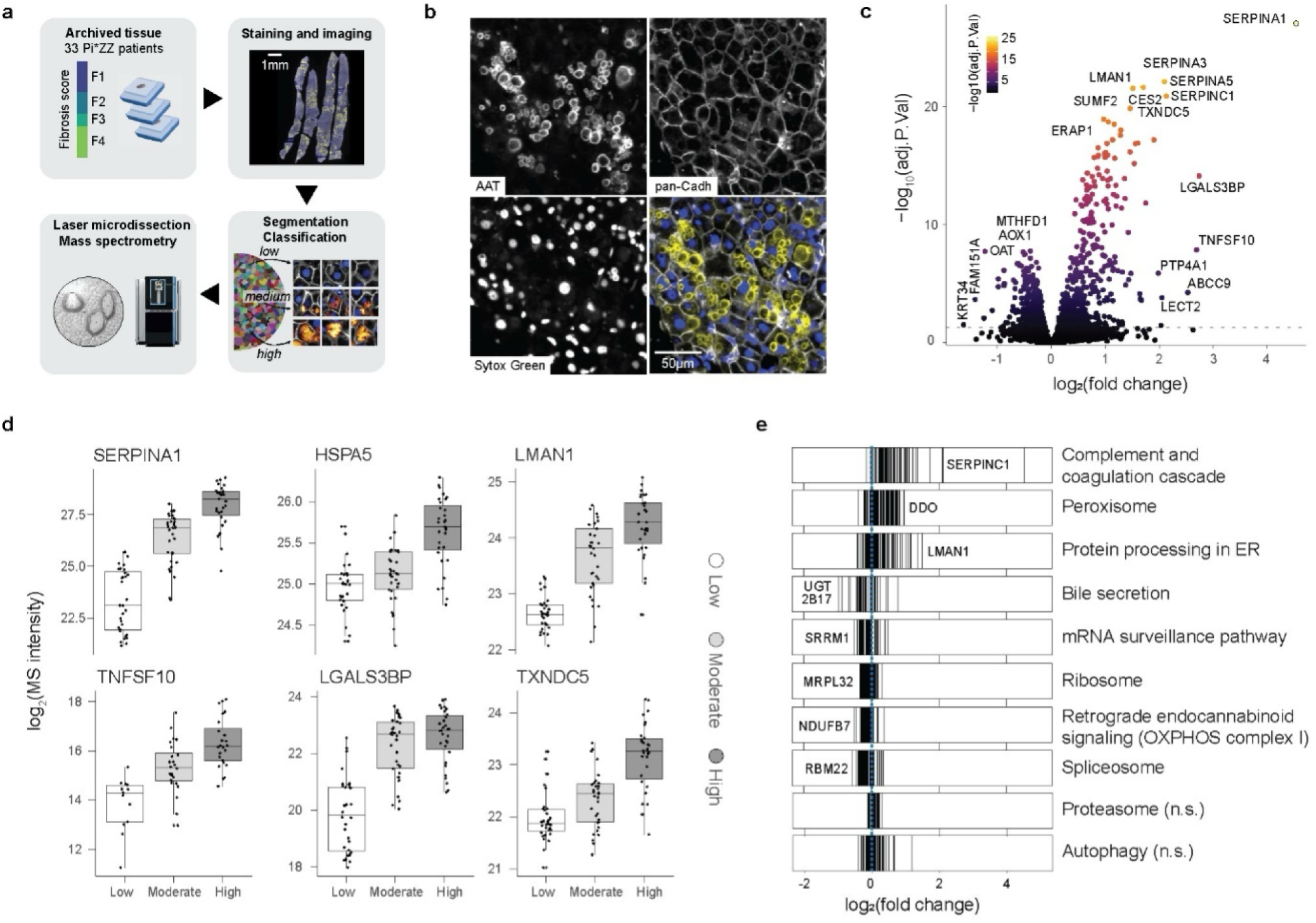
Proteomic mapping of hepatocyte stress response. **a,** Overview of the Deep Visual Proteomics workflow. Fibrosis stages are Kleiner scores. **b,** Immunofluorescence staining of alpha-1 antitrypsin (AAT), the cell outline marker pan-cadherin (pan-Cadh), nucleus (SytoxGreen), and three-color overlay. **c,** Proteomic changes in high versus moderate versus low AAT-accumulating cells. Enriched in high on the right side. Top significant and top changed hits are named (paired two-sided t test with load class as covariable, multiple testing corrected, n = 95 at 100 shapes per sample). **d,** MS intensity of selected proteins across three classes. One dot is one sample from a patient (n = 32). **e,** Significantly (FDR > 0.05) enriched KEGG pathways after Gene Set Enrichment Analysis. Each line is a member of the pathway. n.s. not significant.

Among the most dysregulated hits, we identified other secretory proteins, including many SERPINs, coagulation, and complement factors (Fig. 1c, Extended Data Fig. 1i). This corroborates the notion of ineffective processing and crowding in the ER space, with pathological implications due to the systemic deficiency of multiple plasma proteins^16^. Galectin-3 binding protein LGALS3BP and the apoptotic inducer TNFSF10/TRAIL had the most pronounced positive changes (Fig. 1c and 1d). LGALS3BP is a hepatocyte-produced protein targeted for secretion that is elevated in plasma from patients with liver disease^22^. Reports describing the immune-modulatory activity of LGALS3BP could explain the involvement of immune cells in AATD liver pathology^13,23,24^.

Pathway enrichment analysis showed a strong elevation of proteins related to the three branches of unfolded protein response (UPR) mediated through ATF6, PERK and IRE1 along with a general upregulation of chaperones, accompanied by a reduction of the transcription and translation machinery. This occurred at the expense of physiological functions such as bile secretion (Fig. 1e). Strikingly, many responses converged into a protective response to reactive oxygen species (ROS) with upregulation of thioredoxins and glutaredoxins, including an atypical increase in the peroxisomal compartment and reduction of mitochondrial complex I (Fig. 1d, Extended Data Fig. 1h to 1m). Proteasomal and autophagy proteins remained largely unchanged, and neither did we detect disturbances of calcium homeostasis (Fig. 1d, Extended Data Fig. 1n).

### Early and late-stage responses to proteotoxic stress

Our experimental design, encompassing three aggregate load classes, should allow us to resolve the stepwise progression of molecular events. To determine the sequence in which molecular responses occur during AAT build-up, we first correlated AAT with other protein levels to identify ‘followers’ that tightly track AAT levels. Proteins of the endoplasmic reticulum were among the top ten hits, many destined for secretion (Fig. 2a, Extended Data Fig. 2a and 2b). This included many structurally similar SERPINs, and the tight tracking of AAT levels suggests that these proteins accumulate in tandem with AAT rather than being co-regulated.

**Figure 2:**
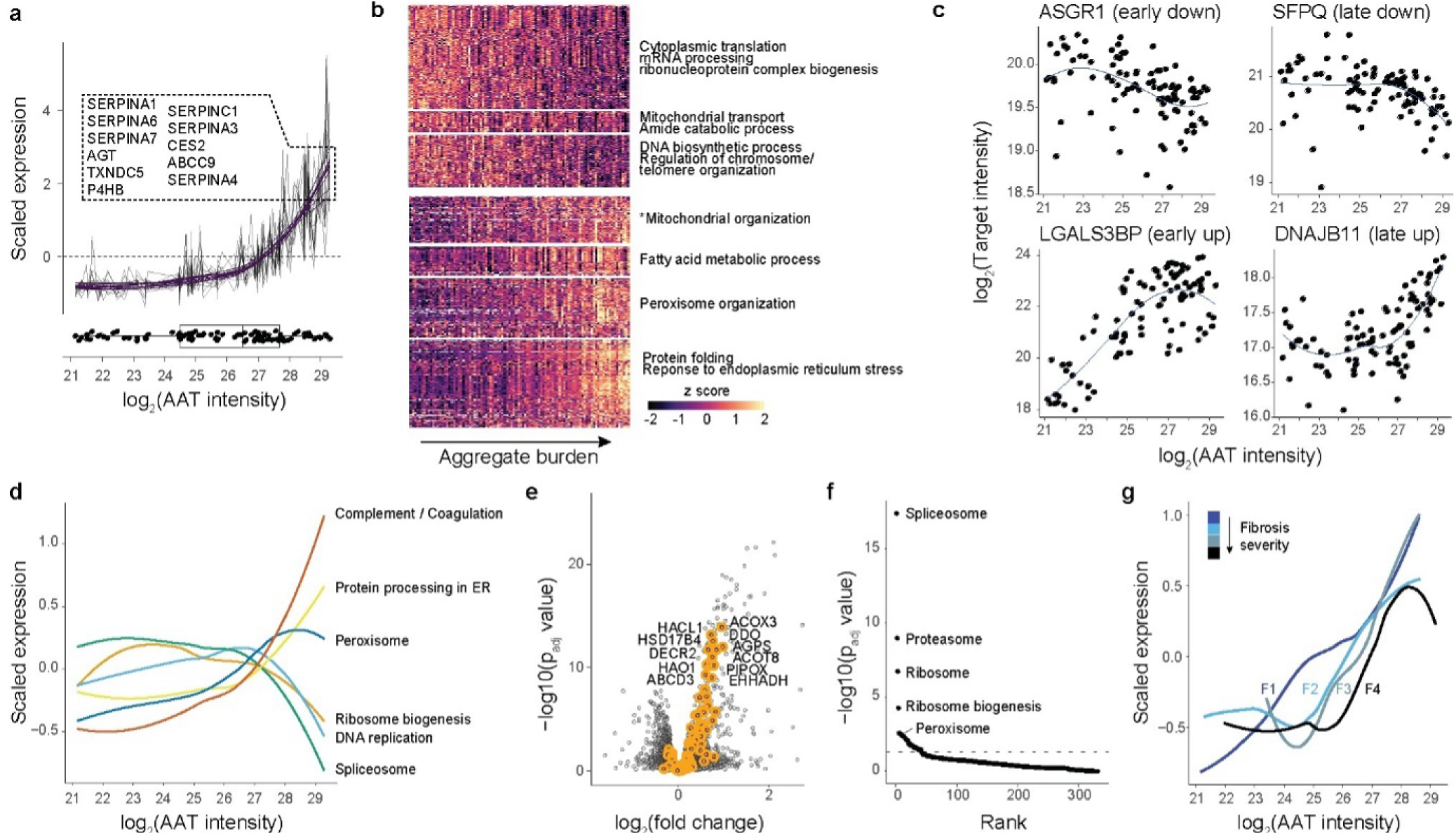
Early and late responses to proteotoxic stress. **a,** Expression profile of the top-ten proteins correlating with AAT. All DVP sample are plotted, and values belonging to the same protein are on one line. Purple, polynomial fit (third order). Boxplot, distribution of AAT expression values along the x axis. **b,** Clustering into early and late responding genes to proteotoxic stress, order on x axis by AAT levels. The y axis was broken into seven groups to achieve good coverage of all response types. Significant KEGG term per box are shown, *not significant. **c,** Pseudo-time expression of top early and late responders by directionality. **d,** Cumulative changes of indicated KEGG pathways expressed as z scores. **e,** Changes of proteins levels across three AAT bins, highlighting peroxisomal proteins. Top significant and top changed hits are named (paired two-sided t test with load class as covariable, multiple testing corrected, n = 95). **f,** Top differential functional categories between F1 and F4 fibrotic samples during early AAT accumulation (log2(AAT intensity) ώ 25; two-sided Wilcoxon test, multiple testing corrected). **g,** Cumulative expression of peroxisomal proteins across four fibrosis stages.

We then categorized proteins into early and late responders to proteotoxic stress caused by AAT accumulation (Fig. 2b). We observed the most consistent relation with AAT load among co-elevated proteins, with the majority (77%) manifesting as late responders and only a smaller fraction as early responders. The immune-modulatory marker LGALS3BP, was most prominent among early responders, followed by the ER cargo receptor MCFD2 together with its co-binder LMAN1 (Fig. 2c). Intriguingly, a strong peroxisomal biogenesis response emerged early on, characterized by the peroxisomal proliferation factor PEX11B and other membrane-integral proteins, along with lipid metabolism and superoxide detoxifying proteins (Fig. 2d and 2e, Extended Data Fig. 2c and 3). In contrast, most proteins of the core machinery of the unfolded protein response appeared later during AAT build-up, despite visual protein accumulation at earlier stages (Fig. 2d, Extended Data Fig. 2d and 2e). The crosstalk between UPR and peroxisomal activity remains poorly understood, yet lipid metabolism, cholesterol metabolism, and ROS detoxification intersect both pathways. Together, the data indicate a dominant increase of the endoplasmic reticulum oxidoreductase 1 alpha (ERO1A), a major peroxide producer (Extended Data Fig. 2b).

We then analyzed samples at various fibrosis stages, revealing major dysregulations with increasing fibrosis stage in proteotoxicity-responsive pathways (Fig. 2f). Notably, this included the peroxisomal response, which showed a gradually prolonged onset time relative to AAT load (Fig. 2g). Importantly, peroxisomal chaperones or chaperone-like proteins remained unaltered, suggesting that peroxisomes are unlikely to contribute to the clearance of unfolded proteins (Extended Data Fig. 2c).

### Single-cell mapping in intact tissue

The accumulation of AAT in intact tissue exhibits a pronounced spatial component. Prior work has demonstrated that AAT accumulates unequally along the zonation gradient from portal to central vein axis in AATD-patients with then Pi*ZZ genotype^13,25,26^. Yet, sharp borders and the absence of gradual changes between neighboring AAT+ and AAT-cells, as well as single positive cells, indicate a more complex picture (Fig. 3a). To map the spatial proteome in these regions, we built upon our previous single-cell DVP workflow^5^ and isolated single shapes from selected regions in 10 µm thick FFPE sections (equivalent to one-third to one-half of a complete hepatocyte) from three F1-stage biopsies. We quantified the proteome of these ‘shapes’ one at a time, allowing us to map back the proteome information onto the tissue with preserved single-cell spatial resolution (Fig. 3a).

**Figure 3:**
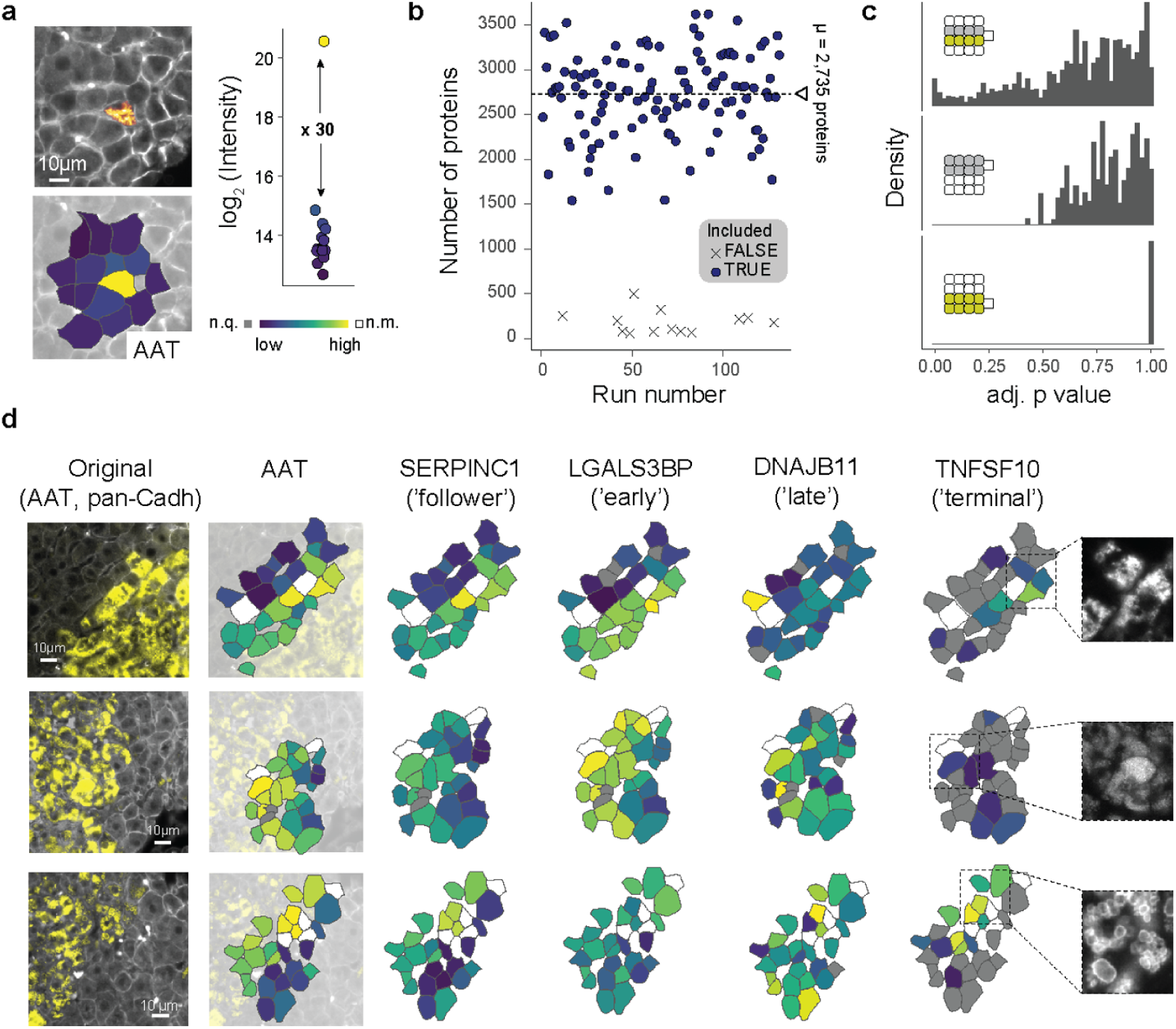
Mapping intact tissue at single cell level. **a,** Enrichment efficiency of the workflow as shown by isolating adjacent cells from FFPE tissue. Proteome quantification of AAT mapped back onto tissue. Boxplot shows AAT expression enrichment. **b,** Number of proteins detected per single shape across all 132 runs. **c,** Distribution of p values when comparing single cells at a border (top, n = 68), direct AAT-neighbours (middle, n = 69) and direct AAT+ neighbours (bottom, n = 49; two-sided unpaired t test after multiple testing correction). **d,** Mapping of proteomic information onto the original microscopic image. Cut-out images show AAT staining only. Gray, protein not quantified (n.q.); white, shape not captured and measured (n.m.) (N = 3, n = 132).

In this way, we quantified the proteome of 132 single shapes in three biopsies at a median depth of 2,735 proteins, and reaching up to 3,600 proteins in some cells (Fig. 3b). The laser capturing proved highly efficient (9.9% dropout rate) and precise, as evidenced by the complete separation of adjacent AAT+ and AAT-cells (Fig. 3a, Extended Data Fig. 4a to 4d). Upon comparing AAT+ and AAT-cells at border regions, we identified similar proteotoxic stress markers as before (Extended Data Fig. 4e to 4g). Interestingly, cells of the first or second row within a border region and within their respective AAT class displayed very similar proteomes (Fig. 3c). Consistent with this, the AAT-accumulation markers LGALS3BP and ERO1A were markedly different between AAT+ and AAT-cells, but not among first and second-order neighbors. Consequently, the data supports an absence of dedicated stress propagation between neighboring cells, suggesting that proteotoxic stress is a cell-intrinsic response.

AAT accumulation has been previously characterized as a peri-portal event^27^. However, our data indicate only partial or no dependence of AAT accumulation on zonation, as evidenced by a drastic change in the expression levels of the portal marker ASS1 at borders, but not HAL and ARG1, or the central markers ADH1 and CYP2E1. Notably, we observed a marked loss of subunits of oxidative phosphorylation in AAT+ cells, including complex IV subunits (mt-CO2, COX5B, COX6C, and others), a signal that was largely undetectable when comparing bulk samples of three groups (Extended Data Fig. 4c, 4h). Importantly, we did not observe any zonation effect in single AAT+ cells compared to AAT-direct neighbors (Extended Data Fig. 4i).

Upon mapping early- and late-responder markers back onto tissue, we found the expected pattern at border regions for SERPINC1 and LGALS3BP, which mirrored AAT levels early on. The late marker DNAJB11 remained unchanged in two of the three samples, indicating that we captured the accumulation event at an early to medium stage (Fig. 3d). However, we detected upregulation of the apoptotic inducer TNFSF10 in the border cells in one sample. Further inspection revealed that the aggregate morphology was markedly different, with a globular phenotype in contrast to amorphous AAT accumulation in the other two samples. Differential expression analysis highlighted intracellular sequestration of iron (FTH1, FTL), the apoptotic marker TNFSF10, and MBL2, as well as several enzymes related to detoxification functions.

### Globular aggregates mark apoptotic cells

Motivated by this observation, we enhanced our DVP workflow to connect cellular phenotypes with proteomic data acquisition. We obtained liver resection samples containing thousands of cells with various AAT aggregate morphologies on one slide. After staining and confocal imaging of 3 µm thick sections of four biological and five technical samples, we segmented cells and transformed the AAT channel signal within cell boundaries into 2048 features representing AAT morphology using the ConvNeXt convolutional neural network^28^. We projected these representations into a two-dimensional space using UMAP and determined 50 equally distributed center points across the image information layer, from which selected the 50 closest cells. These were isolated by laser microdissection and measured by MS, resulting in 250 morphology classes representing a total of 12,500 cells (Fig. 4a).

**Figure 4:**
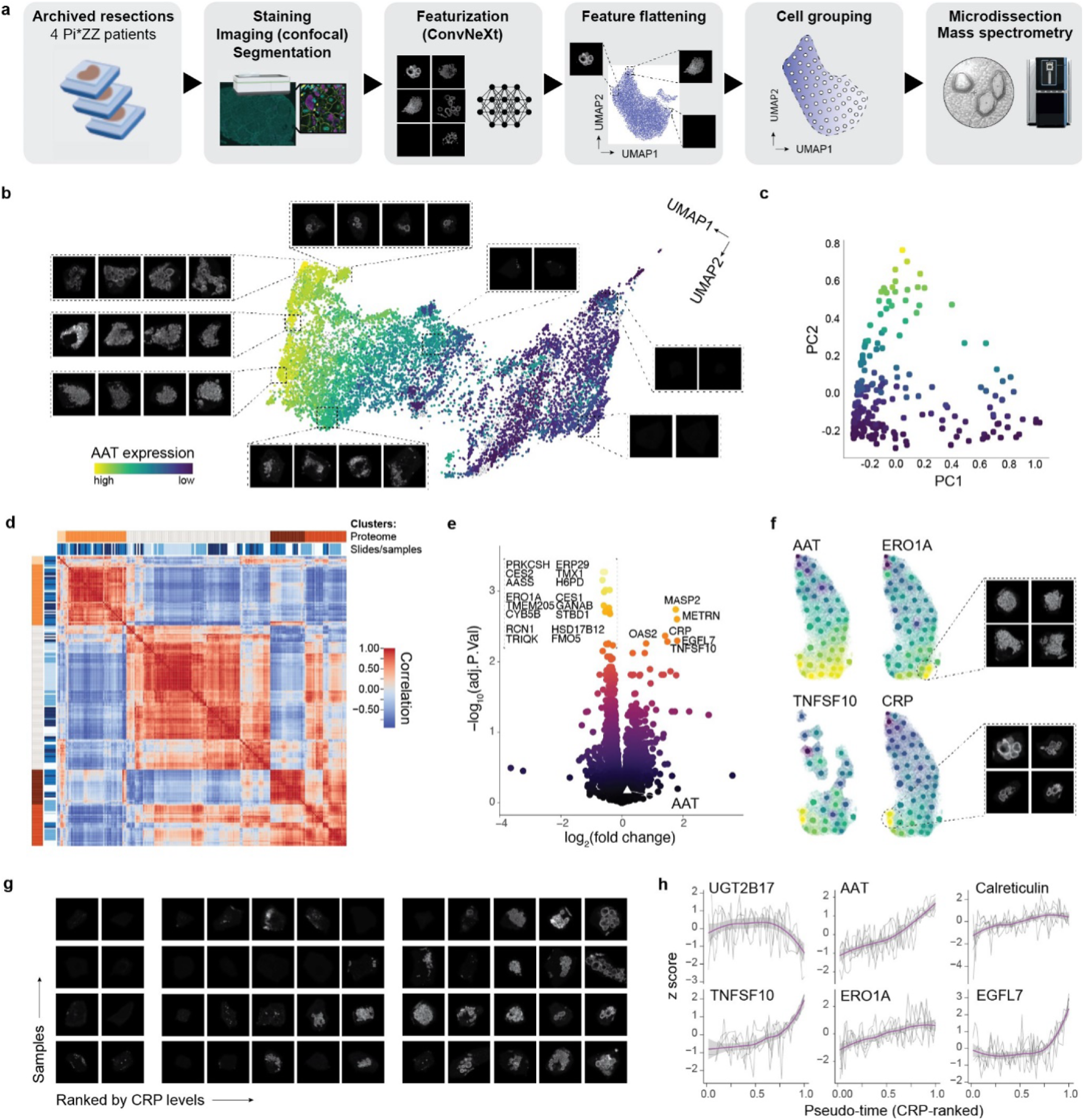
The proteome of cells with various aggregate morphologies (n = 4). **a,** Overview of the CNN-DVP pipeline. **b,** Projection of all laser microdissected cells (12,500) and representative AAT images in indicated areas. Color scheme refers to AAT expression level (proteomic). **c,** Proteomic data of 209s samples (after filtering) reduced by PCA. **d,** Proteomic sample correlation heatmap, indicating proteome clusters based on k means clustering (5 groups manually chosen) and samples slides. **e,** Comparison of proteomes from cells with globular versus amorphous aggregates after selecting for similar AAT levels (AAT indicated as white triangle). Up in globular on the right, top hits annotated (paired two-sided t test after multiple testing correction). **f,** Projection of proteomics data onto image-based UMAP space of one representative sample, with representative images of indicated clusters. **g,** Pseudo time-sorted images of all four biological replicates. Groups mark inflection points of CRP. **h,** Expression levels of indicated proteins in CRP-ranked pseudo-time. Each line is one sample, smoothing curve in purple with 95%-confidence interval in grey.

Employing UMAP to project the representation of these micro-dissected cells into a 2D space validated that the utilized CNN could indeed stratify cells by aggregate morphologies, with aggregate-devoid cells clustering on one end and globular and amorphous morphologies located at the opposite side and clearly separated from one another (Fig. 4b). We achieved a median proteomic depth of 5,970 proteins from the equivalent of 5 to 10 complete hepatocytes (Extended Data Fig. 5a). The main drivers of our proteomic data were dynamic changes in keratins and AAT levels on principal components 1 and 2, respectively (Fig. 4c, Extended Data Fig. 5b to 5d). When grouping samples by proteome into clusters, patient samples were equally distributed across proteomic clusters without apparent genotypic or technical biases (Fig. 4d). As an inverse proof-of-principle, we successfully mapped the proteomic clusters back onto the UMAP image space with clear dimensional separation (Extended Data Fig. 5e). Consistently, samples of one proteome cluster also exhibited the shortest distances to one another on a proteomic UMAP and t-SNE plot (Extended Data Fig. 5f and 5g).

To better understand the molecular responses underlying morphology types, we comparatively analyzed samples with clear globular versus amorphous aggregates (Fig. 4e). Contrary to expectation, markers that typically follow AAT levels, like CES2 and ERO1A, were decreased in globular types. Conversely, the apoptotic inducer TNFSF10 and the inflammatory marker C-reactive protein (CRP) were positively enriched, indicating this to be a terminal phenotype preceding intrinsic or extrinsic apoptosis. We then mapped levels of marker proteins back onto the UMAP-derived image space. Intriguingly, ERO1A and TNFSF10 were localized in two distinct cell populations (Fig. 4f). While ERO1A, indicative of an ongoing UPR response, was highly enriched in amorphous aggregate types, TNFSF10 was mostly present in cells with globular aggregates alongside innate immune system activators. In line with this, Gene Set Enrichment Analysis further identified processes related to cell death as upregulated in globular types (Extended Data Fig. 5h).

Given a rather linear response rate of CRP across the image UMAP space (Fig. 4f), we then sorted all samples in pseudo-time by CRP expression levels. Across all four biological samples, we observed the emergence and disappearance of small corpuscular aggregates despite retained CRP signal. This was followed by a fulminant amorphous aggregation prior to condensation into globular aggregates as a terminal feature before cell death and clearance (Fig. 4g). In addition to TNFSF10, we identified EGF-like domain-containing protein 7 (EGFL7) as a viable marker of this stage that appeared late in the AATD phenotype. Notably, EGFL7 is also upregulated in hepatocellular carcinoma, and high expression levels are associated with poor prognosis^29^. However, a potential link between globular phenotypes and HCC incidence in AATD remains unexplored. This terminal phenotype was further characterized by a stagnating or even declining unfolded protein response in late stages, as evidenced by Calreticulin and ERO1A levels, while reclining levels of proteins such as UGT2B17 suggest the termination of physiological functions in this hepatocyte subtype (Fig. 4h).

## Discussion

We present a pseudo-time resolved proteome of individual hepatocytes undergoing proteotoxic stress due to AAT aggregation. Our findings, derived from FFPE biopsies and resections from patients, provide novel insights into the progression and hepatic manifestation in AAT deficiency. While there are several model systems in the field, including murine models^30^ and patient-derived induced pluripotent stem cells (iPSCs)^31^, our approach uniquely captures responses to proteotoxic stress directly in patients via human tissue specimens representing the full disease spectrum (stages F1-F4). Notably, our data reveal that existing Pi*ZZ models do not accurately recapitulate the UPR, which manifests as a late but fulminant mode of action in our patient-derived samples^1,32^. This discrepancy extends to the globular phenotype, which we now identify as the terminal cellular feature preceding cell death^14^. Our approach strikingly underlines the power of harnessing patient cohorts and tissues. As many potentially druggable targets and pathways are intrinsically more difficult to validate when appropriate model systems are not in place, this inverts the traditional biomedical discovery cycle.

We here developed a single-cell proteomics approach to generate high-resolution maps of adjacent hepatocytes in intact tissue, leveraging recent advancements in ultra-low input mass spectrometry^6,7,33^. Building upon our previous work mapping zonation profiles in frozen mouse liver sections at single-cell resolution^5^, we now quantify 50% more proteins and apply single-cell Deep Visual Proteomics (scDVP) to formalin-fixed tissue. This compatibility with FFPE tissue specimens, the gold standard in diagnostic pathology, expands access to cohorts of virtually any origin, age, and size^34^, broadening the potential applications of this technology.

Our findings indicate that cells without aggregates are not directly affected or triggered by seeding-like mechanisms from adjacent aggregate-bearing cells. However, the presence of large patches of positive cells implies a propagation mechanism. Given the extensive metabolic perturbations observed, including alterations in fatty acid metabolism and detoxification pathways, AAT aggregate formation in one cell may lead to changes in the metabolic microenvironment, thereby inducing stress and proteostatic imbalance in adjacent cells. This hypothesis aligns with other reports in the AATD field and similar mechanisms have been proposed in the context of neurodegenerative proteotoxic disorders where, however, it remains subject of ongoing debate^35,36^.

We present an integration of image featurization and DVP that enables characterization of the entire proteomic and phenotypic lifecycle of stressed hepatocytes in a proteotoxic and fibrogenic liver disease. This methodology establishes a robust framework for dissecting complex cellular processes *in situ* across a spectrum of proteotoxic diseases. This strategy, an example of digital pathology with quantitative and very deep proteomic readout, yielded exceptionally deep proteomes of 6,000 quantified proteins, sufficient to inter most of the functional proteome of a given cell type. Importantly, our datasets are large enough to generate robust models capable of predicting the proteome of a cell based solely on its phenotype. This advancement paves the way for whole-slide proteomics in the future, representing a leap forward in our ability to comprehensively analyze tissue types at exceptional molecular and spatial resolution by mass spectrometry.

The methods developed here recapitulate known disease progression markers while identifying hundreds of additional dysregulated proteins. The present study is necessarily limited in functional followups, yet these novel candidates clearly offer a valuable resource for biological and clinical validation. Of particular clinical relevance, we uncover an early upregulation of the peroxisomal compartment in samples from patients with low-grade liver fibrosis. This response is significantly delayed in high-grade fibrotic samples, suggesting a potential window for therapeutic intervention. PPAR-*α* agonists, such as fibrates, which increase peroxisome load in the liver, may be promising candidates for treating patients with late-diagnosed advanced liver fibrosis due to AATD. Given their well-established safety profiles, we suggest that these drugs could be repurposed for AATD, potentially transforming the treatment landscape of this proteotoxic disorder.

## Methods

### Clinical cohorts and sample preparation

Patient biopsies and explant samples were obtained at two different sites, Odense University Hospital (OUH, Denmark) and Aachen RWTH University Hospital (UKA, Germany). Following ethical guidelines, the clinical data provided here is de-identified by only reporting sample type, fibrosis score, and site of origin.

### OUH patient recruitment

Patients were recruited through the Danish patient organization (Alfa-1 Denmark) and clinical departments for liver and lung diseases as part of a cohort study. The cohort was designed to investigate liver health among non-pregnant adults (minimum age 18 years) diagnosed with AATD of any genotype and carrier status. This specific study includes 16 individuals diagnosed with Pi*ZZ who consented to undergo the procedure. The study was approved by the Danish Ethical Committee (S-2016987), and participants gave informed consent prior to enrollment. Participants without a history of liver transplant or decompensated cirrhosis were offered a percutaneous liver biopsy. The patients underwent liver core needle biopsies at Odense University Hospital (OUH) between 2017 and 2021. Liver core needle biopsies were taken during this period, stored in 4% formalin, and embedded in paraffin. For the assessment of fibrosis stage, FFPE blocks were cut on a microtome into 3µm thin sections and mounted on FLEX IHC slides (Dako, Glostrup, Denmark). Tissue sections were deparaffinized with xylene, rehydrated in serial dilutions of ethanol, and stained with Sirius Red. A certified hepatopathologist (S.D.) assessed the Kleiner fibrosis stage (0-4) according to the Pathology Committee of the NASH Clinical Research Network (NAS-CRN).

### UKA patient recruitment

The recruitment of patients is described in detail in reference^37^. Of this cohort, the present study includes 19 individuals diagnosed with Pi*ZZ, of whom 14 underwent liver core needle biopsies due to medical indication and five received a liver transplantation due to end-stage liver disease. Samples were stored in 4% formalin and embedded in paraffin. Fibrosis stage was assessed after trichrome staining of 5µm thin sections by a certified hepatopathologist. Blocks were stored at room temperature. Ethical approval was provided by the institutional review board of Aachen University (EK 173/15). All participants provided written informed consent and were treated following the ethical guidelines of the Helsinki Declaration (Hong Kong Amendment) as well as Good Clinical Practice (European guidelines).

### Staining

Two micrometer PEN membrane slides (MicroDissect GmbH) were exposed to UV light (254 nm) for one hour and then coated with Vectabond (Vector Laboratories; SP-1800-7) according to the manufacturer’s protocol. Three (DVP, ML) or ten (scDVP) micrometer thin FFPE sections were mounted onto these slides and dried at 37°C overnight. Slides were stored at 4°C until further processing, upon which slides were baked at 55°C for 40 minutes, and then deparaffinized and rehydrated (xylene 2 *×* 2 min, 100% EtOH 2 *×* 1 min, 90% EtOH 2 *×* 1 min, 75% EtOH 2 *×* 1 min, 30% EtOH 2 *×* 1 min, ddH_2_O 2 *×* 1 min). Slides were transferred to prewarmed glycerol-supplemented antigen retrieval buffer (DAKO pH 9 S2367 + 10% Glycerol) at 88°C for 20 minutes, followed by a 20-minute cooldown at room temperature (RT 22°C). After washing in water, sections were blocked with 5% BSA in PBS for one hour, followed by an overnight incubation with primary antibodies in 1% BSA/PBS at 4°C in a humid staining chamber (1:200 mouse IgG1 monoclonal AAT 2C1, Hycult HM2289; 1:200 rabbit recombinant anti-pan cadherin [EPR1792Y], Abcam ab51034). After three washes in PBS for two minutes each, secondary antibodies (1:400 goat anti-mouse IgG1, Invitrogen A21127; 1:400 goat anti-rabbit AF647, Invitrogen A21245) in 1% BSA/PBS were applied for 90 minutes, followed by two 2-minute washes in PBS, 15 minutes in SYTOX™ Green (1:40,000 in PBS, Invitrogen S 7020), and three final 2-minute washes in PBS. Excess liquid was removed and samples were coverslipped using SlowFade Diamond Antifade Mountant (Invitrogen, S36963).

### Imaging

#### Widefield Imaging

For DVP and scDVP experiments (Figures 1–3), sections were imaged using a Zeiss Axioscan 7. For all excitation wavelengths (504 nm, 577 nm, 653 nm), 50% light source intensity was used. The illumination time was specified on one section and applied to all consecutive samples within one experimental group. Three z-stacks at an interval of 2 µm were recorded with a Plan-Apochromat 20*×*/0.8 M27 objective and an Axiocam 712 camera at 14-bit, with a binning of 1 and a tile overlap of 10%, resulting in a scaling of 0.173 µm *×* 0.173 µm. Multiscene images were then split into single scenes, z-stacks combined into a single plane using extended depth of focus (variance method, standard settings), and stitched on the pan cadherin channel using the proprietary Zeiss Zen Imaging software.

#### Confocal Imaging

For experiments with downstream ML applications (Figure 4), sections were imaged on an PerkinElmer OperaPhenix high-content microscope, controlled with Harmony v4.9 software, at 40*×* magnification and 0.75 numerical aperture, with a binning of 1 and a per tile overlap of 10%. Only one z-plane was recorded, which was manually specified for each slide and channel. The three channels were imaged consecutively after deactivation of simultaneous recording to avoid any leakage between channels.

### Cell selection (BIAS)

Images were imported as .czi files into the Biological Image Analysis Software (BIAS) using the packaged import tool^4^. Within BIAS, images were then retiled to 1024*×*1024 pixels with an overlap of 10%, and empty tiles were excluded from further analyses. Cell outlines were identified based on anti-pan cadherin stains using Cellpose 2.0 with the default cyto2 model^38^. Masks were imported into BIAS, and duplicates, as well as cells touching the borders of a tile (0.1% on each side), were removed. Further filtering was applied to retain cells with a minimum size of 3000 pixels, enriching for the hepatocyte population. For classification based on low, medium, and high aggregate load, the cell populations were divided per sample into five classes using a multilayer perceptron (MLP) with the following parameters: weight scale 0.01, momentum 0.01, maximum iterations 10,000, epsilon 0.0005, and 5 neurons in the hidden layer. Classification was based on the AAT (alpha-1 antitrypsin) maximum, median, and mean intensity within the cell outline mask. No human feedback was provided during this process. The low class was attributed to the cells with the lowest normalized mean intensity, medium to the third highest, and high to the highest normalized mean intensity; the other two intermediate classes were dropped. Reference points were selected based on prominent nuclear and histological features. One hundred cells were randomly picked for excision.

For single shape experiments, three characteristic low-fibrosis samples (all F1) and regions were selected that presented with a clear border-like phenotype (i.e., a row of AAT+ cells in direct neighborhood to AAT-cells) or with single AAT+ cells surrounded by AAT-cells. The cells were selected manually in BIAS, starting from the innermost cell and moving spiral-like to the outermost cell, thus avoiding cross-contamination of consecutively cut material.

### Single-cell image generation

Images were flat-field corrected during image acquisition using the Perkin Elmer Harmony software (v4.9). Stitching of the flat-field corrected image tiles was performed using SPARCStools (https://github.com/MannLabs/SPARCStools). The stitched tile positions were calculated using the anti-pan cadherin stains imaged in the Alexa647 channel as a reference and then transferred to the other image channels. During stitching, the tile overlap was set to 0.1, the filter sigma parameter to 1, and the max shift parameter to 50.

The stitched images were then further processed in the python library SPARCSpy (https://github.com/MannLabs/SPARCSpy). Cell outlines were identified based on the 7*×* downsampled anti-pan cadherin stains using Cellpose 2.0 with the pretrained “cyto” model^38^. Segmentation was performed in a tiled mode with a 100px overlap. After resolving the cell outlines from overlapping regions, the resulting segmentation mask was upscaled to the original input dimensions during which the edges of the masks were smoothened by applying an erosion and dilation operation with a kernel size of 7.

Then, the generated segmentation mask was used to extract single-cell image datasets with a size of 280px *×* 280px. During extraction, the same single-cell image masks are used to obtain the pixel information from each channel for each cell. The resulting single-cell images were then rescaled to the [0, 1] range while preserving relative signal intensities. The resulting single-cell image datasets were filtered to only contain cells from within manually annotated regions in the tissue section containing hepatocytes but not fibrotic tissue.

### Cell selection (CNN)

The filtered single-cell image datasets produced by SPARCSpy were further filtered to remove any cells that fell outside the 5 to 97.5% size percentile. Representations of the remaining cells were generated by featurization using the natural image-pretrained ConvNext model^28^. For this, the single-cell images depicting the Alpha-1 channel were rescaled to the expected image dimensions of Npx *×* Npx and triplicated to generate a pseudo rgb image. Inference was then performed using the huggingface transformers package v. 4.26^39^.

The resulting 2048 image features were projected into a two-dimensional space using the UMAP algorithm^40^. The UMAP dimensions were calculated on the basis of the first 50 principal components and the 15 nearest neighbours. Using the spectral clustering algorithm from scikit-learn^41^, the resulting UMAP space was split into 50 clusters. The geometric centre of each cluster was calculated and the 50 cells with the smallest Euclidean distance to the cluster centre were selected for laser microdissection.

Contour outlines of the selected cells were generated in SPARCSpy using the py-lmd package^42^, whereby the cell outlines were dilated with a kernel size of 3 and a smoothing filter of 25 was applied. Furthermore, the number of points defining each shape were compressed by a factor of 30 to improve LMD cutting performance. The cutting path, i.e. which cell is cut after one another, was optimized using the Hilbert algorithm (https://github.com/galtay/hilbertcurve).

### Laser microdissection

After aligning the reference points, contour outlines were imported, and shapes were cut using the LMD7 (Leica) laser microdissection system in a semi-automated mode with the following settings: power 45, aperture 1, speed 40, middle pulse count 1, final pulse 0, head current 42-50%, pulse frequency 2,982, and offset 190. The microscope was operated with the LMD beta 10 software, calibrated for the gravitational stage shift into 384-well plates (Eppendorf 0030129547), leaving the outermost rows and columns empty. To prevent sorting errors, a ‘wind shield’ plate was placed on top of the sample stage. Plates were then sealed, centrifuged at 1,000 g for 5 minutes, and subsequently frozen at *−*20°C for further processing.

### Peptide preparation and Evotip loading

Peptides were prepared as previously described using a BRAVO pipetting robot (Agilent) as per reference^43^. Briefly, 384-well plates were thawed, and shapes (both combined and individual) were rinsed from the walls into the bottom of the well with 28µL of 100% acetonitrile (ACN). The wells were completely dried in a SpeedVac at 45°C, followed by the addition of 6µL of 60mM triethylammonium bicarbonate (TEAB, Supelco 18597) (pH 8.5) supplemented with 0.013% n-Dodecyl-beta-D-maltoside (DDM, Sigma-Aldrich D5172). Plates were sealed and incubated at 95°C for one hour. After adjusting to 10% ACN, samples were incubated again at 75°C for one hour. Subsequently, 6ng and 4ng of trypsin and Lys-C protease, respectively, in 1 µL of 60 mM TEAB buffer were added to each sample, and proteins were digested for 16 hours at 37°C. The reaction was quenched by adding trifluoroacetic acid (TFA) to a final concentration of 1%. Peptide samples were then frozen at *−*20°C.

For loading, new Evotips were first soaked in 1-propanol for one minute, then rinsed twice with 50 µL of buffer B (ACN with 0.1% formic acid). After another 1-propanol soaking step for three minutes, the tips were equilibrated with two washes of 50 µL buffer A (0.1% formic acid). Samples were loaded into 70 µL of pre-loaded buffer A. Following one additional buffer A wash, the peptide-containing C18 disk was overlaid with 150µL buffer A and briefly centrifuged through the disk. All centrifugation steps were performed at 700g for one minute. The final tips were stored in buffer A for a maximum of four days prior to LC-MS.

### LC-MS data acquisition

The peptide samples were analyzed using an Evosep One liquid chromatography (LC) system (Evosep) coupled to an Orbitrap Astral mass spectrometer (Thermo Fisher Scientific). Peptides were eluted from the Evotips with up to 35% acetonitrile (ACN) and separated using an Evosep low-flow “Whisper” gradient for DVP samples, or an experimental Evosep “Whisper Zoom” gradient for single shapes and DVP-ML samples, with a throughput of 40 samples per day (SPD) on an Aurora Elite TS column of 15 cm length, 75 µm internal diameter (i.d.), packed with 1.7 µm C18 beads (IonOpticks). The column temperature was maintained at 50°C using a column heater (IonOpticks).

The Orbitrap Astral mass spectrometer was equipped with a FAIMS Pro interface and an EASY-Spray source (both Thermo Fisher Scientific). A FAIMS compensation voltage of *−*40V and a total carrier gas flow of 3.5 L/min were used. An electrospray voltage of 1900V was applied for ionization, and the RF level was set to 40. Orbitrap MS1 spectra were acquired from 380 to 980 m/z at a resolution of 240,000 (at m/z 200) with a normalized automated gain control (AGC) target of 500% and a maximum injection time of 100 ms.

For the Astral MS/MS scans in data-independent acquisition (DIA) mode, we experimentally determined the optimal methods across the precursor selection range of 380-980 m/z: (a) For DVP samples, a window width of 5 Th, a maximum injection time of 10 ms, and a normalized AGC target of 800% were used. (b) For DVP-ML samples, a window width of 6 Th, a maximum injection time of 13 ms, and a normalized AGC target of 500% were applied. (c) For single shapes and other DIA scans, the window width was optimized based on precursor density across the selection range of 380-980 m/z. A total of 45 variable-width DIA windows were acquired with a maximum injection time of 28 ms and an AGC target of 800%. The isolated ions were fragmented using higher-energy collisional dissociation (HCD) with 25% normalized collision energy.

Detailed method descriptions will be provided in a default format with each supplementary data table upon publication.

### Spectral searches and normalization

The raw files were searched together with match-between run in library-free mode within each experimental group with DIA-NN v1.8.1^44^. A FASTA file containing only canonical sequences was obtained from Uniprot (20,404 entries, downloaded on 2023-01-02), and the disease-causing amino acid was manually changed (E342K). We allowed a missed cleavage rate of up to 1, and set mass accuracy to 8, MS1 accuracy to 4, and the scan window to 6. Proteins were inferred based on genes, and the neural network classifier was set to ‘single-pass mode’. For DVP and DVP-ML samples, precursor intensities in the ‘report.tsv’ file were then normalized using the directLFQ GUI at standard settings including a minimum number of non-nan ion intensities required to derive a protein intensity of one^45^. The single shape data was additionally median normalized to a set of proteins quantified across all samples (621 proteins quantified in 100% of included samples), thereby correcting for the dependence of protein numbers on shape size^5^.

### Data analysis and statistics

Data was analyzed using R version 4.4.1. The directLFQ output file ‘pg matrix.tsv’ was utilized for all subsequent data analysis, including the reported protein counts. Samples were included if the number of protein groups exceeded the mean minus (a) 1.5 standard deviations for DVP and single shape samples, resulting in 1.0% (1/96) and 10.6% (14/132) dropouts, respectively; and (b) 0.5 standard deviations for DVP-ML samples, resulting in 16.4% (41/250) dropouts. This lower cutoff was selected after manual inspection of the data distribution. Although some samples were collected in technical duplicates per patient biopsy, only the first replicate was used for statistical analyses and all reported measurements were taken from distinct samples. Coefficients of variation were calculated on non-transformed intensity values. For principal component analysis (PCA), the R package PCAtools 2.16.0 was used on a complete data matrix, removing the lower 10% of variables based on variance. Statistical analyses were performed assuming normality using the limma package version 3.60.3 with two-sided moderated t-tests and “fdr” as a multiple testing correction method. A per-patient statistical pairing was applied for DVP and single shape experiments. Intensity and fold changes are reported as log2-transformed values unless indicated otherwise. Gene Set Enrichment Analysis (GSEA) was conducted using WebGestalt 2024 against the indicated databases, with a false discovery rate (FDR) of < 0.05 considered significant^46^. Interaction networks were calculated with STRING database at standard settings^47^. The timing of responses ranked by the absolute difference between B values of limma’s moderated t test comparing three AAT load groups: low to moderate, and moderate to high. Only proteins that were significant in either or both comparisons were considered. Differential pathway expression across fibrosis stages was calculated by fitting a linear model through log2-transformed intensity values of individual proteins in samples with log2(AAT)-intensity < 25, and the slopes of proteins in a particular pathway were compared between F1 and F4 samples by a two-sided Wilcoxon rank test without assumption of normality. Indicated p values are corrected for multiple testing using the ‘fdr’ method. Spatial data was mapped using the ‘simple features’ package.

## Data availability

The mass spectrometry proteomics data will be made available upon publication.

## Acknowledgements

We thank our colleagues at the Department of Proteomics and Signal Transduction at the Max Planck Institute of Biochemistry as well as our colleagues at the Center for Proteome Research in Copenhagen for their input and support. We are particularly grateful for the technical assistance of Dirk Wischnewski, and for input from Thierry Nordmann, Marvin Thielert and Vincenth Brennsteiner. We thank the Computing Centre and the Imaging Facility of the MPI of Biochemistry for their support and resources. F.A.R. is an EMBO postdoctoral fellow (ALTF 399-2021). S.C.M. is a PhD fellow of the Boehringer Ingelheim Fonds. This study has been supported by the Horizon-2020 under the MICROB-PREDICT program (M.M., A.K., no. 825694); by the Max Planck Society for Advancement of Science (M.M.); by a grant from the Alpha-1 Foundation (F.A.R.) and Alfa-1 Denmark (A.K.); by the Deutsche Forschungsgemeinschaft DFG through SFB 1382 (P.S., ID 403224013); P.S. holds a Heisenberg professorship (STR1095/6-1).

## Contributions

Conceptualization: F.A.R., K.H.T., S.C.M., P.S. and M.M. Project teams were led by S.C.M. (image analysis and machine learning), K.H.T. (clinical data), and S.S. (single-cell analysis). Methodology: F.A.R., S.C.M., S.S. Software: S.C.M., M.L., N.A.S. Validation: F.A.R., C.A.M.W., M.O., M.W., M.Z., J.K. Formal Analysis: F.A.R., S.C.M., M.L. Investigation: F.A.R., S.C.M., K.H.T., S.S., M.L., C.A.M.W., M.W., M.Z., J.K. Resources: K.H.T., M.F., S.D., P.B., A.W., O.F., S.F., A.K., P.S., M.M. Data Curation: F.A.R., S.C.M. Writing – Original Draft: F.A.R., M.M. Writing – Review & Editing: all authors. Visualization: F.A.R., S.C.M., M.L., Supervision: F.A.R, P.S., M.M.

---

**Extended Data Fig. 1.**
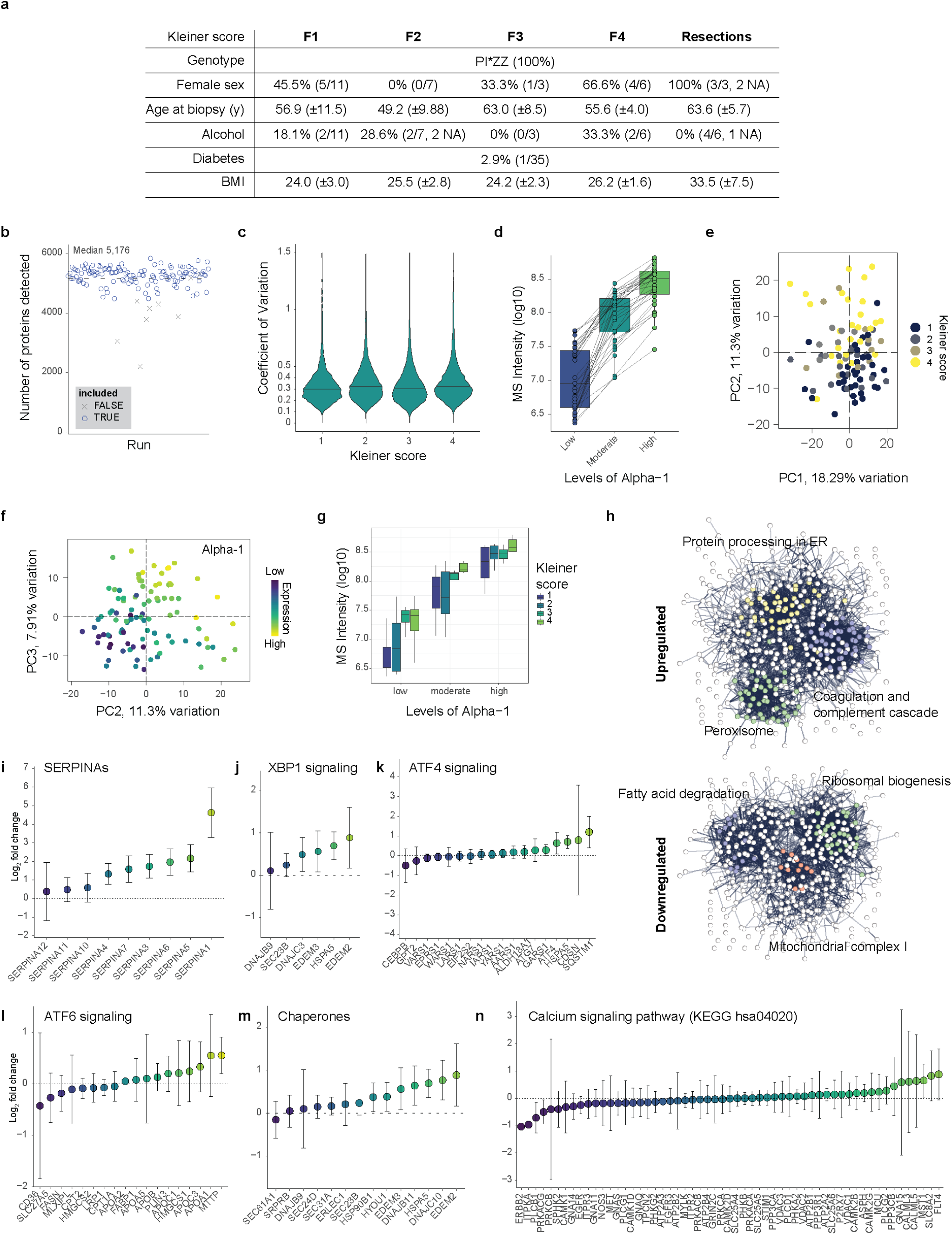
Proteomic mapping of hepatocyte stress response (Part 1). **a,** Summary of clinical metadata expressed in number of patients, or percentages with absolute numbers in brackets. Mean ± SD. **Proteomic mapping of hepatocyte stress response (Part 2). b,** Number of proteins detected across all runs prior to exclusion of technical replicates (n = 134). Upper dotted line: median number of protein groups. Lower dotted line: Median – 1.5 SD. Samples below were excluded and are marked as a cross. **c,** Coefficient of variation across fibrosis stages. **d,** MS intensity of alpha-1 antitrypsin in the three distinctly microdissected cell classes. **e,** Principal component analysis with principal components 1 and 2 color by fibrosis stage, and **f,** with principal component 2 and 3 colored by alpha-1 antitrypsin level. Each dot is one sample (n = 95). **g,** Levels of alpha-1 antitrypsin by fibrosis stage across the three microdissected cell classes (n = 32 patients). **h,** STRING interaction network of significantly (FDR < 0.05) upregulated (top) or downregulated proteins in cells (see Fig. 1c). **i – n,** levels of selected proteins in indicated pathways in cells with compared to without aggregates. Circles indicate mean, bars are SD across patient samples (n = 32). The proteins in i to m were manually selected, n is retrieved from KEGG.

**Extended Data Fig. 2.**
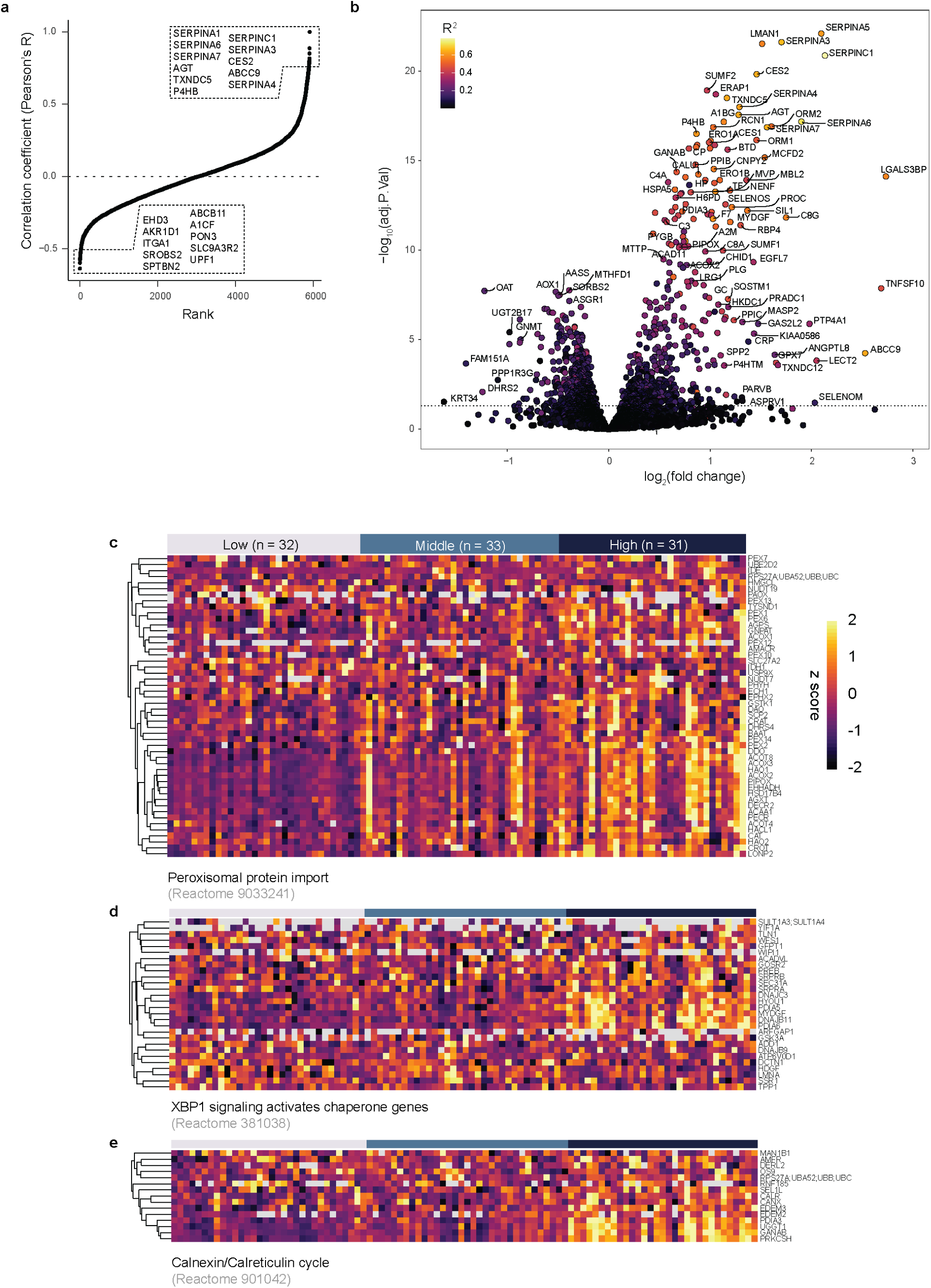
Early and late responses to proteotoxic stress (Part 1). **a,** Pearson’s R correlation coefficient of each detected protein with alpha-1 antitrypsin levels calculated per MS sample. Top and low-10 protein names are indicated in boxes. **Early and late responses to proteotoxic stress (Part 2). b,** Proteomic changes in high versus moderate versus low AAT-accumulating cells colored by their R^2^ value against alpha-1 antitrypsin expression. Enriched in high on the right side. Top significant and top changed hits are named (paired two-sided t test with load class as covariable, multiple testing corrected, n = 95 at 100 shapes per sample). **c,** Expression levels of indicated proteins colored by z score (assuming normality) across all samples split by load class and related to peroxisomal protein import, **d,** XBP1 signaling and **e,** the Calnexin/Calreticulin cycle. Database IDs given below each graph (n = 95 in 32 patients).

**Extended Data Fig. 3.**
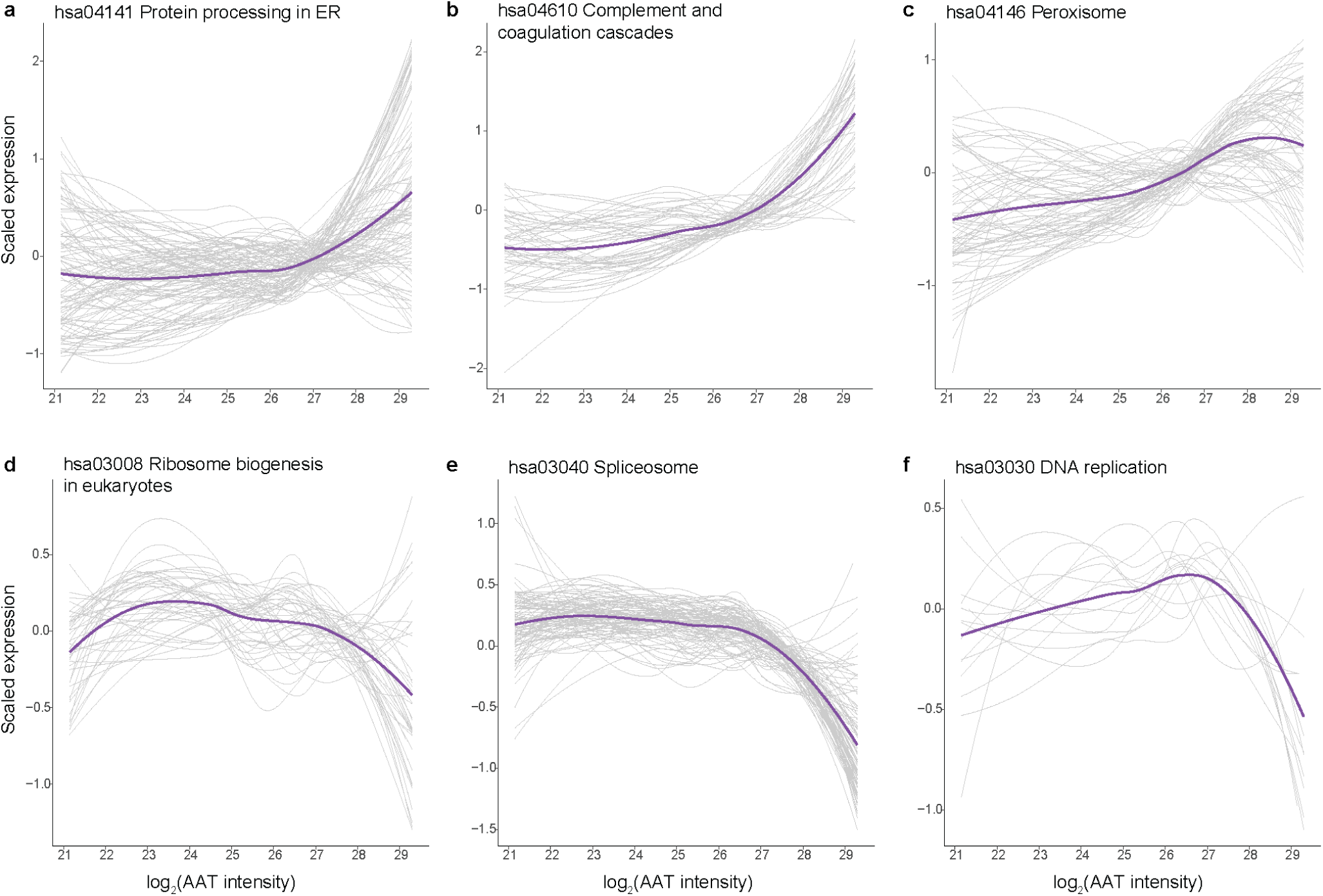
Changes of functional pathways. **a-f,** Scaled intensity (z scored) of all detected proteins in indicated KEGG pathways against AAT intensity. ‘hsa00000’ are KEGG identifiers. Purple line is the local regression (span 0.75, degree 2).

**Extended Data Fig. 4.**
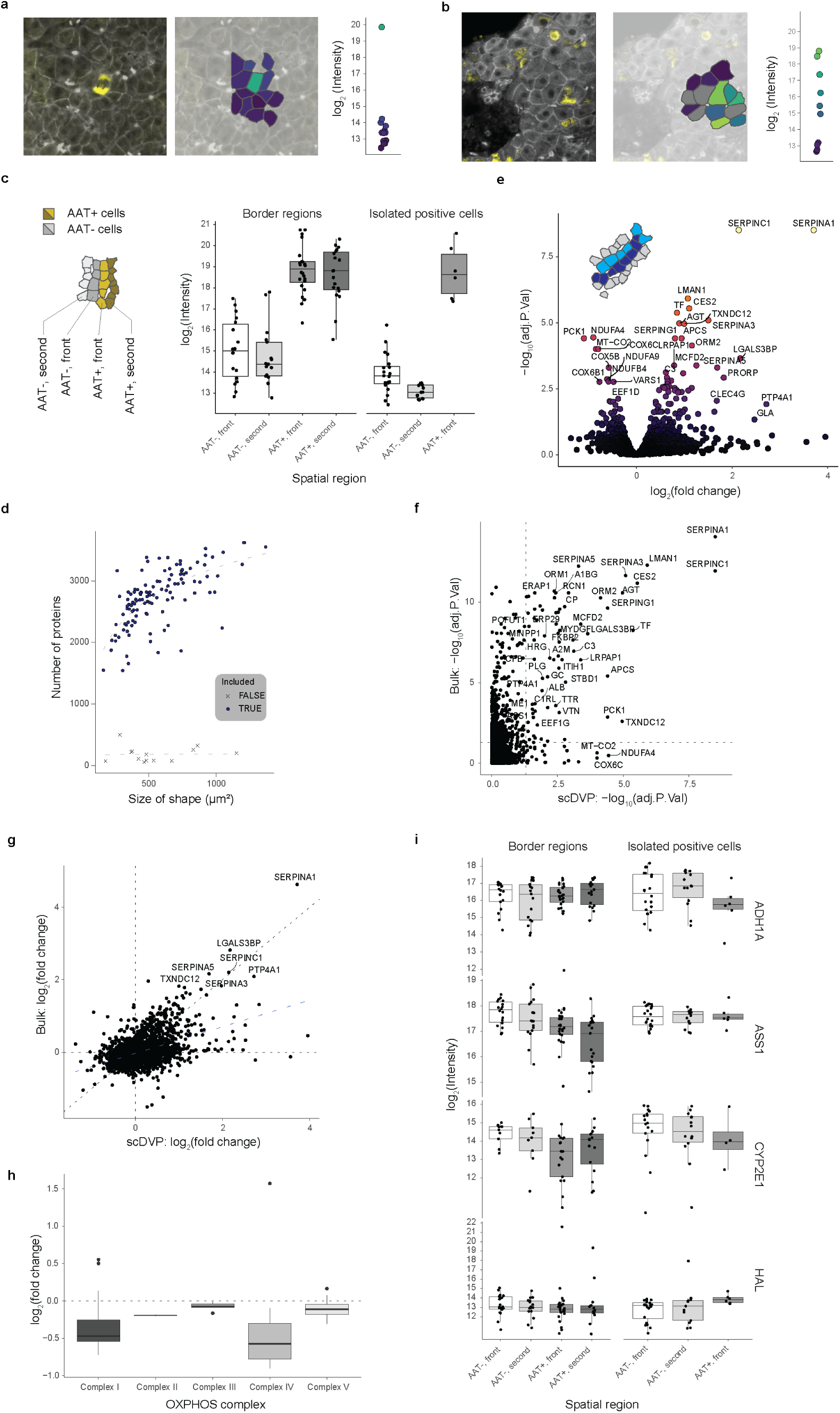
The single-cell proteome (Part 1). **a and b,** Color-coded AAT expression in regions with single-positive cells. AAT expression levels of all indicated shapes are also shown in the dot graph on the right of each spatial mapping **The single-cell proteome (Part 2). c,** Expression of AAT in indicated regions determined by immunofluorescence signal across all included samples (n = 118). **d,** Number of proteins detected in relation to the cut shape area. Excluded samples are indicated with a cross. **e,** Statistical comparison of AAT+ and AAT-cells at the three regions classified as ‘borders’ (paired two-sided t test, multiple testing corrected, 30 AAT+ cells and 38 AAT-cells). **f,** Comparison of adjusted p-values and **g,** log2(fold changes) of AAT+ and AAT-single shape comparisons on the x axis versus cells along the accumulation gradient (refer to Fig. 1 and 2) on the y axis. Statistics as in e, and Fig. 1c. **h,** Relative expression levels of subunits of the oxidative phosphorylation system (OXPHOS) in AAT+ versus AAT-single shapes. Proteins are retrieved from Mitocarta 3.0^49^. **i,** Expression of protein indicated on the right in respective spatial region. Periportal markers: ASS1 and HAL; pericentral markers: ALDH1A1 and CYP2E1. The boxes are first and third quartiles, the thick line is the median, whiskers are ±1.5 interquartile range and outliers are indicated as individual points.

**Extended Data Fig. 5.**
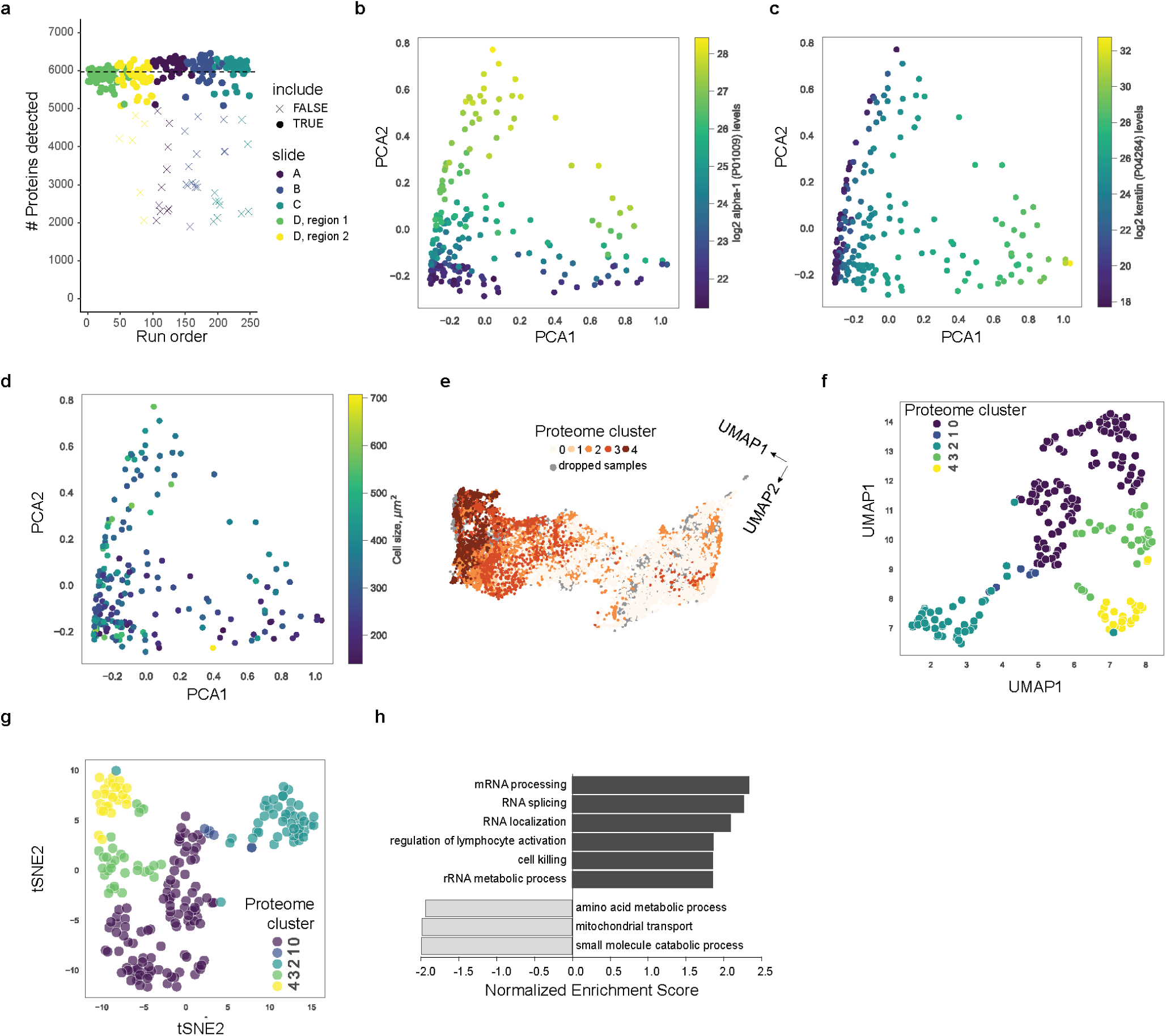
The proteome of cells with various aggregate morphologies. **a,** Number of protein groups detected per sample. Each dot is one sample, the horizontal line indicates the mean across all included samples (n = 209 included, n = 41 excluded and marked with a cross). Exclusion criteria were that the number of detected proteins was smaller than mean minus 0.5 SD. **b,** Principal component analysis of all included samples with AAT, **c,** KRT1 expression levels, or **d,** shape size color coded (n = 209). **e,** Annotation of the proteome cluster in Fig. 4d onto the image space UMAP. Dropped samples are in grey (n = 12,500). **f,** Representation of individual samples color coded by proteome cluster in a proteomic UMAP, or **g,** tSNE space (n = 209). **h,** Gene Set Enrichment Analysis (GO: Biological Process noRedundant) of globular versus amorphous aggregate types.

